# Biodegradable Poly(*β*-amino amide)s Enable Efficient RNA Delivery and Spleen Targeting

**DOI:** 10.1101/2025.03.03.641302

**Authors:** Xuejin Yang, Jingya Xiao, Yajie Zhai, Leqi Liao, Han Qiu, Ye Chen, Daryl Staveness, Xiaoyu Zang

**Affiliations:** N1 Life, Inc., 446 S Hillview Dr, Milpitas, CA 95035, USA; Accuredit Therapeutics, 111 Wusongjiang Avenue,Guoxiang Street, Wuzhong District, Suzhou, Jiangsu 215124, CN

**Keywords:** RNA transfection, gene delivery, gene editing, RNA research, RNA therapeutics, targeted delivery, protein translation, RNA translation

## Abstract

RNA therapeutics are emerging as transformative modalities in clinical applications and have become a key area in life science research. While lipid nanoparticles (LNPs) have become the leading platform for RNA delivery in preventive and therapeutic products, they still face significant challenges in achieving efficient extrahepatic delivery and maintaining long term shelf stability. Here we report the development of a series of biodegradable poly(*β*-amino amide) (PBAA) polymers, detailing their design, synthesis, and performance as gene delivery vehicles both *in vitro* and *in vivo*. These cationic polymers, featuring interspersed trialkylamine motifs, provide a readily tunable functional handle and facilitate complexation with gene cargo. The redox-sensitive disulfide motifs introduce a redox-responsive decomposition pathway for the polymer backbone, triggering cargo release during intracellular delivery. The results herein demonstrate that these polymers offer remarkable efficiency in encapsulating and delivering translation-competent cargos, including mRNA and CRISPR-Cas based gene editing tools. Notably, a dodecyl modified PBAA transporter has achieved over 97% gene editing efficiency *in vitro*, and over 97% spleen-targeting selectivity in a murine model. Additionally, it produces stable nanoparticles that maintain their physicochemical properties at 4 °C for up to two weeks without addition of excipients such as PEG, offering a cost-effective solution for RNA therapeutics supply chains. As the transformative impact of RNA and other nucleic acids continues to build within the pharmaceutical industry and beyond, it is imperative that the supporting technologies evolve in stride to maximize said impact. The tunable and biodegradable PBAA polymer designs presented herein are illustrative examples of how high-level functional performance can be acheived in conjunction with the critical targeting, formulation, and operational simplicity needed of state-of-the-art transfection and delivery technologies.

## Introduction

The emergence of RNA-based therapeutics represents a paradigm shift in modern medicine, offering unprecedented opportunities to address previously untreatable genetic disorders, cancers, and infectious diseases.^[1,2]^ Following the remarkable success of mRNA vaccines during the COVID-19 pandemic,^[3,4]^ which demonstrated the clinical potential of RNA therapeutics at a global scale, there has been renewed interest in expanding the application of RNA-based medicines across diverse therapeutic areas. The approval of patisiran (Onpattro, 2018), givosiran (Givlaari, 2019), lumasiran (Oxlumo, 2020), and inclisiran (Leqvio, 2021) also presents RNA interference as a valid therapeutic approach to various genetic disorders.^[5]^ Unlike traditional small-molecule drugs or protein therapeutics, RNA therapeutics can potentially target any gene of interest, enabling precise manipulation of disease-relevant pathways through various mechanisms including protein expression, gene silencing, or gene editing.

However, the therapeutic potential of RNA molecules has yet to be fully unlocked due to limitations of delivery technologies.^[6]^ RNA is inherently unstable, susceptible to enzymatic degradation, and carries large numbers of negative charges that impede cellular uptake. Moreover, RNA molecules must overcome multiple biological barriers to reach their intended site of action, including but not limited to: avoiding immune recognition, achieving tissue-specific targeting, facilitating cellular internalization, and enabling endosomal escape.^[7]^ The complexity of these delivery challenges has largely restricted the clinical application of RNA therapeutics.

While lipid nanoparticles (LNPs) represent the most clinically advanced delivery platform, their preferential accumulation in the liver has limited their utility for targeting other tissues and organs. This tissue tropism, combined with potential toxicity concerns and manufacturing complexities, has motivated the development of alternative delivery strategies.

The comprehensive understanding of both the promises and challenges of RNA therapeutics underscores the importance of developing novel delivery technologies that can expand the therapeutic window and tissue targeting capabilities of RNA-based medicines. Synthetic carriers that can achieve efficient extrahepatic delivery while maintaining favorable safety profiles remain a critical unmet need in the field. Recent advances in polymer chemistry and materials science have opened new opportunities for designing delivery vehicles with enhanced tissue specificity, improved cellular uptake, and controlled release properties.^[8–10]^

Cationic polymers, including synthetic types such as poly(ethyleneimine) (PEI)^[11]^ and poly(amidoamine) (PAMAM) dendrimers,^[12]^ have been extensively used in nucleic acid delivery owing to their strong cationic properties. However, their excess cationic charge and lack of biodegradable bonds are associated with poor biocompatibility and toxicity, limiting their clinical applications.^[8]^ To address these challenges, research has focused on developing synthetic polymers with improved biodegradability while optimizing their structural tunability and functional diversity to enhance their delivery performance. For instance, charge-altering releasable transporters (CARTs), synthesized via ring-opening polymerization, possess unique self-immolative degradation mechanisms.^[13,14]^ These CART polymers have demonstrated efficient delivery and transfection of mRNA and siRNA into cells and animals. By modifying lipid types in the side chains, adjusting lipid spacing, and tailoring the degree of polymerization for each block, the organ tropism and cell selectivity of CARTs can be fine-tuned.^[14–18]^ Poly(*β*-amino ester) (PBAE)^[19]^ and poly(*β*-amino amide) PBAA^[20,21]^ polymers, which can be easily synthesized through Michael addition of amines into acrylates, have also been widely explored. PBAEs, modified in their polymer topologies, including linear and highly branched, as well as through structural variations in their backbone, side chains, and chain-end capping groups, have demonstrated the ability to tune the delivery efficiency and organ selectivity of RNA and DNA.^[19,22–26]^ In comparison, PBAAs, bearing amide linkages, are inherently more stable against hydrolysis than the ester bonds in PBAEs. This stability provides opportunities to enhance material durability while maintaining flexibility for tunable and controllable degradation, achieved by incorporating degradable linkages into the backbone, such as bio-reducible disulfide bonds^[27–29]^ and photosensitive moieties.^[30]^ However, PBAAs remain less extensively investigated than the PBAE counterpart, and PBAA-centric structure-activity relationships (SARs) remain underexplored.

The work presented herein showcases the advantages available to PBAA-based polymers, demonstrating the ease and scalability of polymer synthesis, high versatility in structural modifications, inherent stability, and tunable biodegradability. Furthermore, establishing SARs, with a particular focus on transfection efficiency and organ tropism for systemic RNA delivery, is pivotal to advancing their clinical applications.

In this study, a combinatorial library of 36 PBAAs incorporating bio-reducible disulfide bonds in the backbone was designed, synthesized, and investigated to establish foundational PBAA SARs in nucleic acid delivery. Key parameters, including polymer composition (homopolymer vs. random copolymer), electronic effects (hydrophobic vs. hydrophilic, saturated vs. unsaturated), steric effects (linear vs. branched), and the length of functional groups in the side chains and end groups were systematically evaluated both *in vitro* and *in vivo*. The top-performing selected PBAA achieves stable polyplex formulation, enables efficient *in vitro* gene editing, and demonstrated over 97% spleen targeting, making it a promising candidate for effective therapeutic applications.

## Results and Discussion

### 1. Design and synthesis of PBAA polymer carriers

The poly(*β*-amino amide) (PBAA) polymers were synthesized via a two-step Michael addition reaction between primary amines and bis(acrylates), resulting in either homopolymers or random copolymers (**Figure 1A** and **1B**). In the first step, the polymer backbone was formed, followed by the addition of an end-capping reagent in the second step to terminate any free acrylate chain ends. If Boc protecting groups were present on the side chains or end groups, they were removed in a subsequent deprotection step (Step 3) prior to testing.

**Figure 1.**
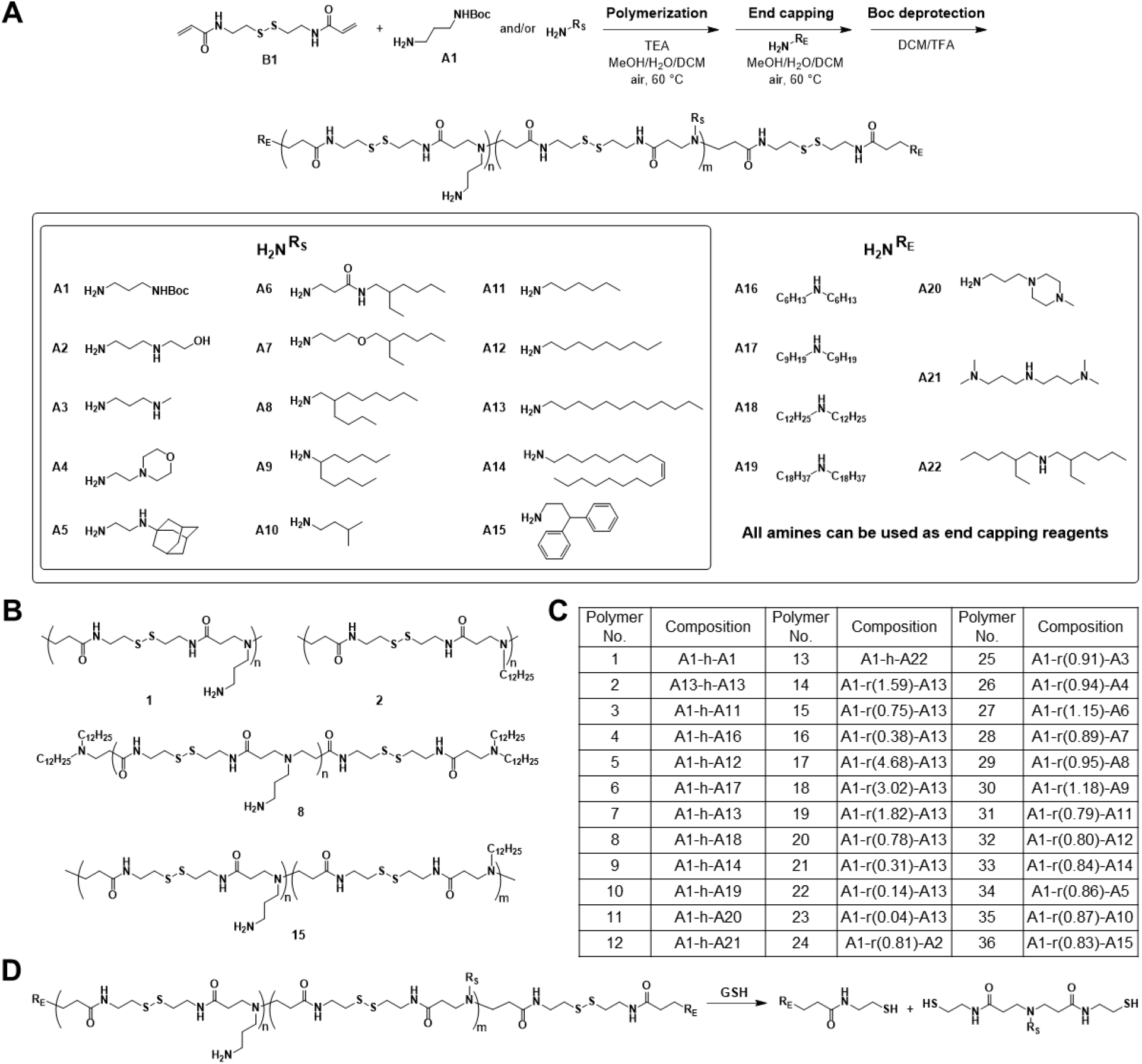
(A) The reaction scheme of biodegradable PBAAs and the chemical structures of amine building blocks, with **R**_**S**_ groups functioning as side chains and **R**_**E**_ groups functioning as end capping units. (B) Examples of homopolymers and random copolymers. (C) The table shows polymer carriers and their compositions. Homopolymers were named **A1-h-AX**, where **AX** represents the amine used for end capping, while random copolymers were named **A1-r(lipid content)-AX**. (D) Polymer degradation in a reducing environment.

Disulfide bonds within the polymer backbone, introduced from the *N,N*′-Bis(acryloyl)cystamine monomer (**B1** in **Figure 1A**), enable rapid and environmentally responsive degradation in the reducing environment of the cytosol. With this molecular degradation mechanism, these polymers break down into segments or small molecules, as illustrated in **Figure 1D**, which accelerates the decomplexation with nucleic acids.^[31,32]^ The amine building blocks employed in the polymerization or the chain-end capping step included cationic, lipophilic, or aryl groups, as shown in **Figure 1A**. In this study, most homopolymers were synthesized using *N*-Boc-1,3-propanediamine (**A1**) as the side chain, paired with various end groups (**AX**). Random copolymers were generated when two different amines were loaded simultaneously during polymerization, with **A1** commonly serving as one of the repeating units. The end-capping step provided an additional opportunity for library diversification, though in this study, the end-capping amine was typically the same as the second amine monomer (**AX**) introduced during polymerization.

The polymers were characterized by GPC and NMR. The molecular weight of homopolymers remained relatively consistent (3 to 5 kDa) before and after chain-end capping. The lipid content of the random copolymers, defined as the ratio of the N-Boc group from **A1** to the second amine group **AX** (approximately m/n), was estimated through NMR analysis. Homopolymers were named as **A1-h-AX**, where **AX** represents the amine used for end capping, while random copolymers were named **A1-r(lipid content)-AX**.

### 2. *In vitro* evaluation of eGFP mRNA delivery and transfection using PBAA polymer carriers

The performance of PBAA polymers in eGFP mRNA delivery was evaluated using HEK 293T cell line. All polymers were dissolved in RNase-free water and prepared as stock solutions at a concentration of 2.5 mg/mL. Nanoparticles were formulated at an N/P ratio of 25 with an eGFP mRNA dose of 40 ng/well.

As shown in **Figure 2**, homopolymers capped with lipophilic end groups, such as **Polymer No. 5** (nonyl), **7** (dodecyl), and **9** (oleyl), containing C_9_–C_18_ alkyl chains, demonstrated higher mRNA transfection efficiency and expression levels compared to those with hydrophilic end groups, including **Polymer No. 1** (1,3-propanediamine), **11** (piperazine), and **12** (dimethylpropylamine). Notably, additional alkyl chains at the end group further improved the ability of mRNA delivery. For instance, the polymer modified with didodecylamine (**Polymer No. 8**) at the chain end exhibited substantially higher transfection efficiency than its counterpart capped with dodecylamine (**Polymer No. 7**). Polymers with shorter C_6_ and C_9_ alkyl groups (**Polymer No. 4** cf. **3, Polymer No. 6** cf. **5**) showed less pronounced improvements. Interestingly, increasing the alkyl chain length to C_18_ (**Polymer No. 9** and **10**) did not enhance efficacy further, suggesting an optimal chain length for maximizing performance.

**Figure 2.**
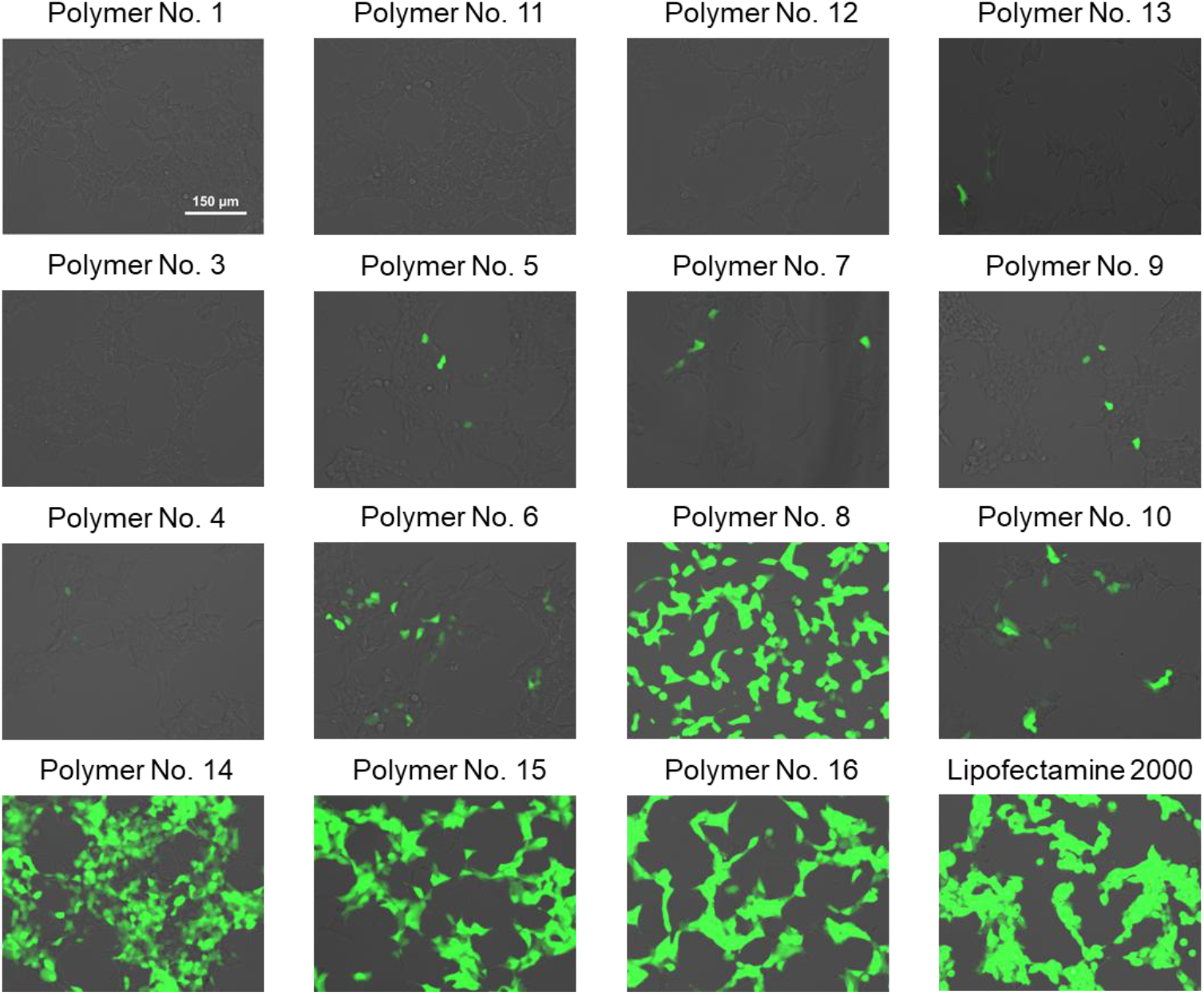
eGFP expression of HEK 293T cells after treating with PBAA/eGFP mRNA nanocomplex for 24 hours. Fluorescence microscope images were captured to assess the performance of selected PBAA polymers (Scale bar, 150 μm).

Compared to the homopolymer end-capped with dodecylamine (**Polymer No. 7**), dodecylamine random copolymers with significantly higher lipid content (lower NH2/C12 ratio), such as **Polymer No. 14** (1.59), **15** (0.75), and **16** (0.38), exhibited a substantial improvement in eGFP mRNA expression levels.

These results clearly demonstrated that polymers with hydrophobic lipids attached to chain ends (chain-end modifications) and dangling from the polymer backbone (polymers having hydrophobic lipid side chains) can induce more efficient delivery of mRNA into cells. Additionally, increasing hydrophobic lipid content in the polymer, as compared to the end-capping strategy is somewhat more effective at improving transfection efficiency. Notably, random copolymers, with enhanced tunability of lipid content, emerge as a more promising system for exploration compared to chain-end modified homopolymers.

### 3. Expanding the PBAA random copolymer library and screening *in vitro*

Based on the *in vitro* screening results above, we focused our synthesis efforts on expanding the random copolymer library. To identify the optimal lipid content for efficient mRNA delivery, additional dodecylamine random copolymers with varying lipid contents (ranging from 0.04 to 4.68) were synthesized and screened *in vitro* (**Figure S1**).

The performance of these polymers in eGFP mRNA delivery was evaluated in HEK 293T cells, with the commercially available JetMESSENGER serving as the positive control. As shown in **Figure S1**, the GFP signal strengthened with an increase in hydrophobic lipid content (corresponding to a decrease in NH_2_/C_12_ ratio from 4.68 to 0.14). However, further increase in lipid content resulted in a decline in the fluorescence signal.

Flow cytometry was used to further quantify these findings. As illustrated in **Figure 3A**, mean fluorescence intensity (MFI) increased with hydrophobic lipid content during the initial 4 hours of transfection, reaching saturation at a lipid content of 0.31 (**Polymer No. 21**), comparable to JetMESSENGER. At 28 hours post-transfection, a peak MFI was observed for **Polymer No. 20** (lipid content 0.78), with both **Polymer No. 20** and **21** showing significantly higher MFI compared to JetMESSENGER. The trends for transfection efficiency followed a similar pattern at both 4 and 28 hours (**Figure 3B**). Notably, **Polymer No. 20, 21**, and **22** achieved over 92% transfection efficiency at 28 hours post-transfection, outperforming JetMESSENGER.

**Figure 3.**
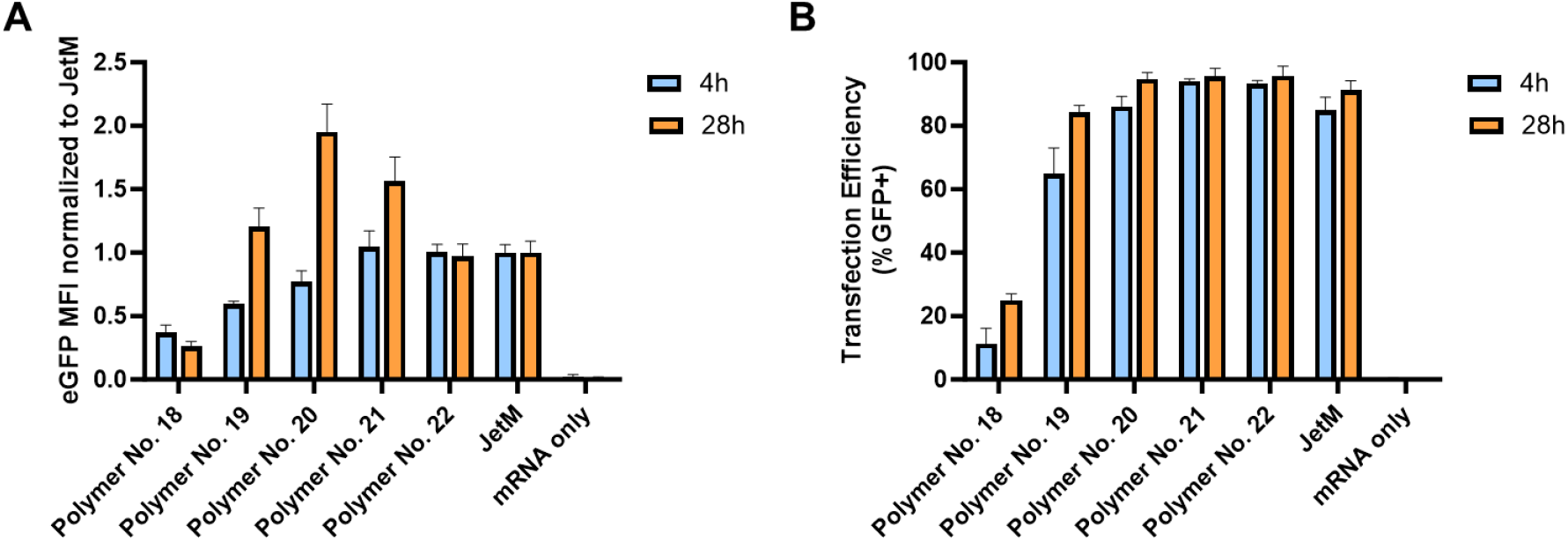
Lipid content screening of dodecyl random copolymers in HEK 293T cells was performed. (A) MFI and (B) the percentage of GFP-positive cells were analyzed 24 hours post-transfection using flow cytometry. Results are shown as mean ± SD.

Based on these findings, we expanded the random copolymer library by synthesizing polymers with diverse side chains while maintaining a lipid content similar to **Polymer No. 20** (0.78), which could be achieved using a 1:1 loading ratio of two different amines.

The performance of these newly synthesized polymers was assessed through Fluc mRNA transfection studies in HEK 293T cells. As shown in **Figure 4**, random copolymers incorporating more hydrophobic lipids (e.g., those with longer hydrocarbon chains and/or alkene moieties) demonstrated higher luciferase mRNA expression levels. For example, among polymers with similar lipid content, the oleyl random copolymer (**Polymer No. 33**) produced a stronger luminescence signal compared to the dodecyl (**Polymer No. 15**), nonyl (**Polymer No. 32**), and isoamyl (**Polymer No. 35**) random copolymers. In contrast, random copolymers with more hydrophilic lipids, such as **Polymer No. 24, 25**, and **26**, exhibited negligible luminescence signals.

**Figure 4.**
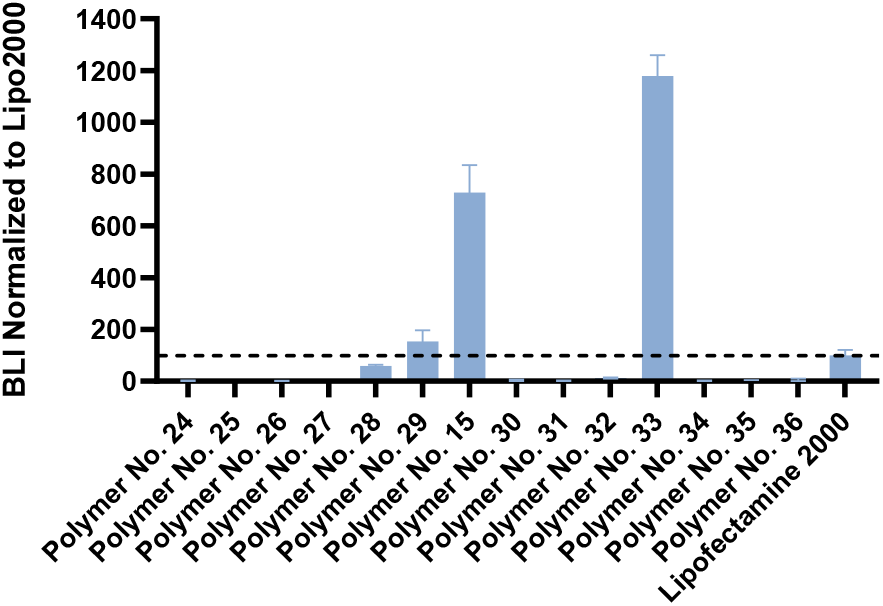
Evaluation of random copolymers with different side chains for luciferase mRNA transfection in HEK 293T cells. Total luminescence signals were normalized against the positive control, Lipofectamine 2000 (with its luminescence signal set at 100).

To further explore the role of side-chain structure on transfection performance, we compared random copolymers with side chains at the same atom count but differing configurations. Interestingly, **Polymer No. 15**, having a linear C_12_ side chain, outperformed **Polymer No. 29**, which has a branched C_12_ side chain.

Additionally, compared to **Polymer No. 29**, which has an all-carbon branched alkyl chain, **Polymer No. 28**, with an oxygen atom incorporated into the side chain, and **Polymer No. 27**, featuring a more electron-withdrawing amide bond, showed progressively lower luminescence signals.

Expanding on the structural diversity, random copolymers with bulky lipids, such as adamantyl (**Polymer No. 34**), and 3,3-diphenylpropyl (**Polymer No. 36**), were also evaluated. However, these copolymers showed minimal transfection signals compared to the positive control group, Lipofectamine 2000, indicating that overly bulky lipids may impede effective mRNA delivery.

Although the oleyl random copolymer (**Polymer No. 33**) exhibited the highest luminescence signals, it faced solubility challenges at stock concentrations of 2.5 mg/mL or higher, limiting its suitability for *in vivo* applications. Consequently, subsequent investigations focused on dodecyl random copolymers, which demonstrated both strong transfection performance and better compatibility in various studies.

### 4. *In vivo* evaluation of Fluc mRNA delivery using PBAA polymer carriers

To further investigate the structure-activity relationship (SAR) identified above, PBAA polymers were evaluated *in vivo* for luciferase mRNA delivery through systemic administration in mice. In the first trial, **Polymer No. 1, 6, 8, 11, 14, 15, 16** were selected and tested. Mice were dosed with nanocomplexes containing 5 µg luciferase mRNA (Fluc mRNA) and imaged after 6 and 16hours post-administration.

**Figure 5A** summarizes the total luminescence observed in BALB/c mice treated with PBAA/Fluc mRNA complexes and the negative control. In most groups, luminescence signals peaked at 6 hours and decreased by several folds at the 16-hour timepoint. Dodecylamine-based random copolymers, such as **Polymer No. 14, 15**, and **16** demonstrated significant total luminescence signals, offering the best performance among all tested trials. Interestingly, while the homopolymer end-capped with didodecylamine (**Polymer No. 8**) was comparably effective *in vitro*, it resulted much lower luminescence *in vivo* relative to the dodecylamine-based random copolymers. Additionally, homopolymers end-capped with shorter lipids, such as the dinonyl group (**Polymer No. 6**), or with hydrophilic end groups, such as 1,3-propanediamine (**Polymer No.1**) and piperazine (**Polymer No. 11**), showed lower or undetectable bioluminescence signals.

**Figure 5.**
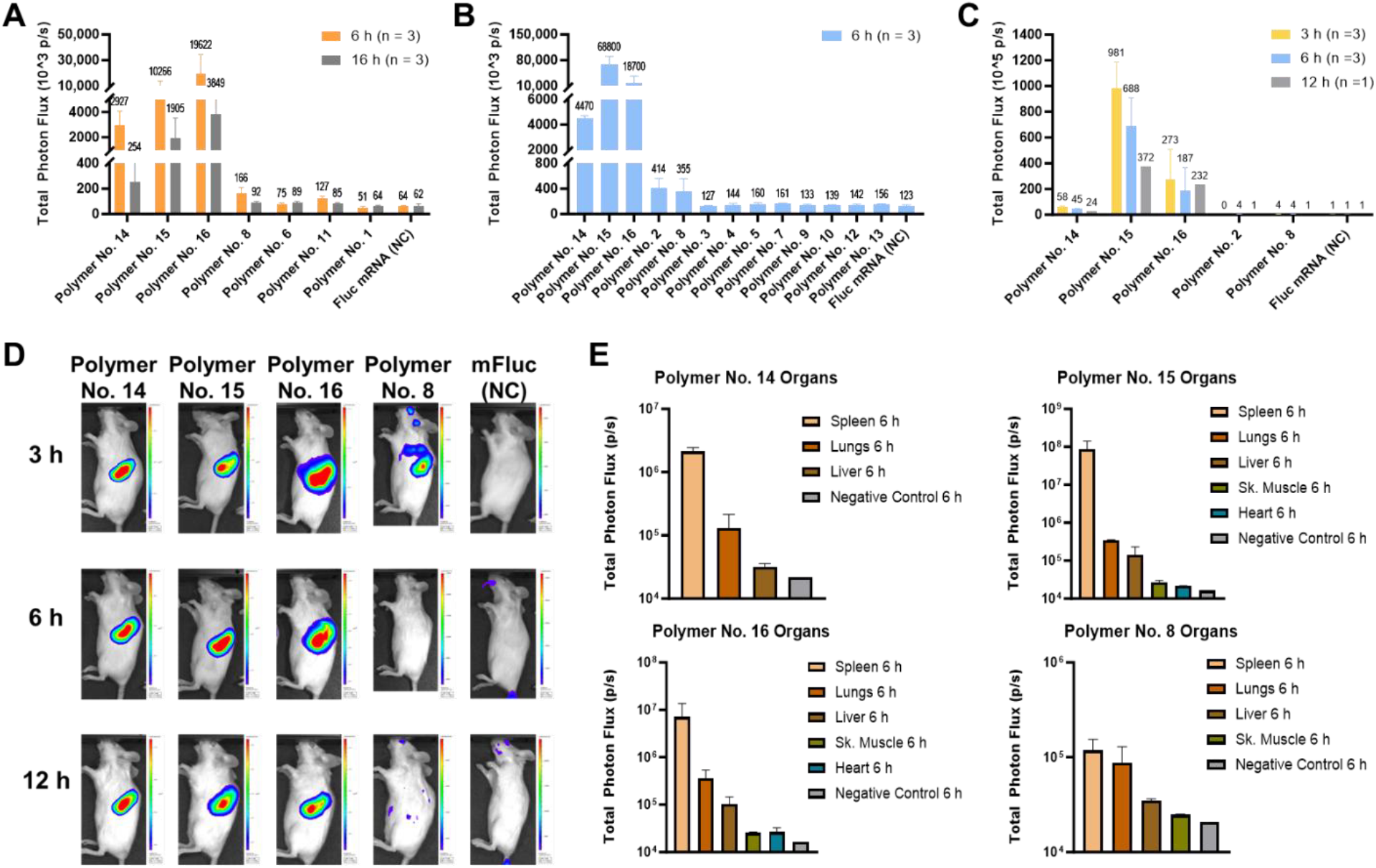
*In vivo* screening of PBAA polymers for Fluc mRNA delivery to mice via intraveneous injection at a dose of 5 µg/mouse. (A) Quantification of bioluminescence imaging (BLI) signals in mice (side view) during the first-round study at 6 h and 16 h. Quantification of BLI signals in mice (side view) at 6 h (B) and comparison of top-performing candidates at various time points (3 h, 6 h, and 12 h) (C) in the second-round study. *In vivo* BLI of mice (D) and BLI signal quantification in excised organs (E) for top-performing candidates in the second-round study (n = 3).

More *in vivo* studies were conducted to further investigate the SAR and assess the biodistribution at the same mRNA dose (5 µg/mouse). Four polymers from the first round of testing (**Polymer No. 8, 14, 15**, and **16**), along with additional homopolymers featuring different end groups, were evaluated (**Figure 5B, 5C**) in the second-round study. The kinetics of mRNA expression *in vivo* were assessed more closely by adding in an earlier time point (t = 3 h). Major organs, including the brain, heart, lungs, liver, spleen, stomach, intestines, kidneys, and skeletal muscle, were immediately harvested and imaged *ex vivo* after mice were sacrificed.

Dodecyl random copolymers (**Polymers No. 14, 15**, and **16**) once again demonstrated significantly higher mRNA delivery efficiency than that of the homopolymers with various end-capping groups (**Polymer No. 3** – **13** as shown in **Figure 5B** and **5C**), measured by luminescence signals. Notably, **Polymer No. 15** exhibited slightly higher luminescence than **Polymer No. 16**, reversing the ranking observed in the first study. As previously observed, the didodecyl end-capped homopolymer (**Polymer No. 8**) produced luminescence signals two orders of magnitude lower than the random copolymer (**Polymer No. 15**). Replacing the end-group modification from didodecyl (**Polymer No. 8**) to hexylamine (**Polymer No. 3**), dihexylamine (**Polymer No. 4**), 2-ethyl-hexyl (**Polymer No. 13**), nonyl (**Polymer No. 5**), dinonylamine (**Polymer No. 6**), oleyl (**Polymer No. 9**), or dioctadecylamine (**Polymer No. 10**) led to reduced delivery efficacy.

Remarkably, dodecyl random copolymers (**Polymer No. 14, 15**, and **16**) exhibited noticeably higher luminescence signal accumulation in the spleen, as shown in **Figure 5D**. And the organ-specific targeting was further validated through *ex vivo* imaging of isolated organs and bioluminescence quantification (**Figure 5E**). These random copolymers delivered Fluc mRNA to multiple organs including spleen, lung, liver, and skeletal muscle, achieving over 90% spleen targeting, and **Polymer No. 15** displayed the highest spleen selectivity. In contrast, the didodecyl-end-capped homopolymer, **Polymer No. 8**, showed approximately 40% spleen selectivity. In the other homopolymer trials, no specific luminescence signals were detected in the spleen, lungs, liver, skeletal muscle, or heart at any time point, with signal levels comparable to those of the negative control group.

A third round of *in vivo* studies was conducted to specifically explore the effects of the N/P ratio and mRNA dose on systemic Fluc mRNA delivery and biodistribution. **Polymers No. 15** and **16**, which exhibited strong performance and spleen tropism in previous studies but ranked differently, were chosen for detailed comparison. PBAA/Fluc mRNA nanocomplexes were formulated by systematically varying the N/P ratio and mRNA dose while maintaining a consistent polymer amount across all formulations (**Table S1**).

Female ICR Mice (6–8 weeks old) were divided into seven groups and received either a vehicle (PBS) or nanoparticles carrying Fluc mRNA via tail vein injection (i.v.), as detailed in **Table S1**. Live imaging (abdominal view) was conducted for each group after 6 hours administration. Mice displaying bioluminescence signals were sacrificed, and the major organs, including spleen, liver, and lungs were dissected and imaged immediately *ex vivo*.

As illustrated in **Figure 6A, Polymer No. 15**/Fluc mRNA at N/P ratio of 5 and mRNA dose of 50 μg (**Table S1**, Group 2) produced the strongest *in vivo* bioluminescence signal among all tested groups. No significant differences were observed between N/P ratios of 10 and 25 for **Polymer No. 15** (**Table S1**, Group 3 cf. Group 4) or **Polymer No. 16** (**Table S1**, Group 6 cf. Group 7) at mRNA dose of 25 µg and 10 µg, respectively.

**Figure 6.**
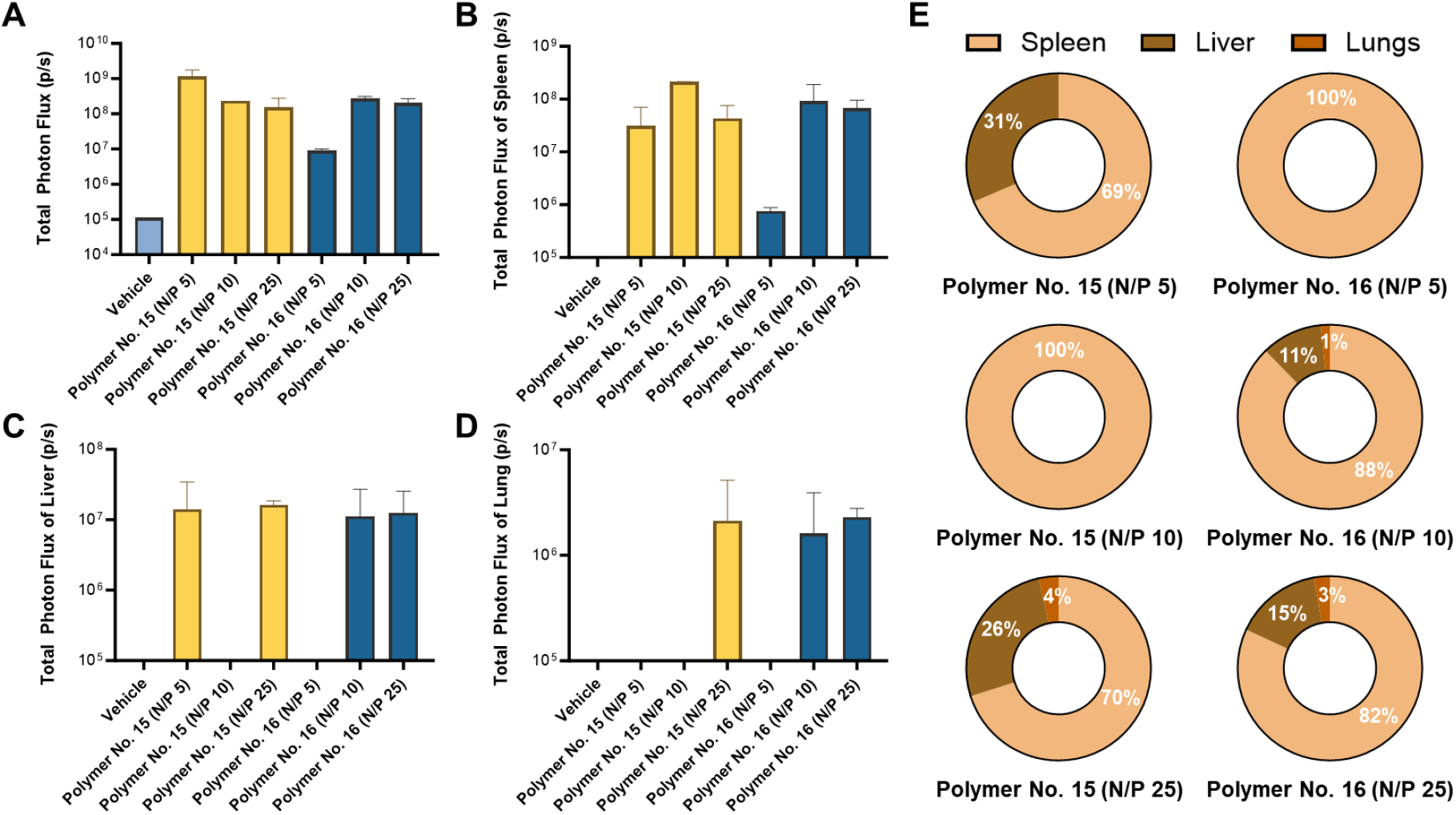
(A) Quantification of *in vivo* bioluminescence in ICR mice dosed with PBAA/Fluc mRNA nanocomplexes at 6 h. Quantification of *ex vivo* bioluminescence in the spleen (B), liver (C), and lungs (D) at 6 h. (E) Percentage distribution of bioluminescence signals among the three major organs. The percentage distribution was calculated by dividing the bioluminescence signal from the spleen, liver, and lungs by the total signal from these organs combined.

**Figure 6B–E** depict the biodistribution of luciferase mRNA expression. Among the three major organs analyzed, bioluminescence signals were predominantly localized in the spleen, followed by the liver and lungs. Both **Polymer No. 15**/Fluc mRNA at N/P ratio of 10 and **Polymer No. 16**/Fluc mRNA at N/P ratio of 5 demonstrated exclusive spleen tropism. However, the bioluminescence intensity for the latter was two orders of magnitude lower than that of the former.

Overall, **Polymer No. 15** and its formulation with Fluc mRNA at an N/P ratio of 10 was determined to be the optimal condition, offering a combination of high total bioluminescence signal and strong spleen selectivity.

### 5. PBAA random copolymer enables gene editing *in vitro*

Given the successful delivery of eGFP and Fluc mRNA *in vitro*, we extended our studies to the delivery of CRISPR/Cas9 based system for gene editing applications. CRISPR/Cas technology enables precise, sequence-specific gene editing and has been rapidly adapted for various applications, including but not limited to the correction of disease-causing mutations, genetically engineering of animals for *in vivo* studies and food production, and advancements in agricultural biotechnology.^[33–35]^

To evaluate the PBAA polymer within this gene-editing context, **Polymer No. 15** was employed to simultaneously deliver both spCas9 mRNA and sgRNA cargos into hepatocyte Huh7 cell line. Various weight ratios of the spCas9 mRNA and sgRNA gene cargos were evaluated (ranging from 0.11 to 39), with a fixed total RNA dose of 10 or 20 ng per well (96-well plate scale). The two RNA cargos were pre-mixed, and the polymer/RNA complexes were formulated at an N/P ratio of 10.

As shown in **Figure 7A, Polymer No. 15** successfully delivered spCas9 mRNA and sgRNA in a single formulation mixture. The system enables gene editing in Huh7 cells across a broad range of spCas9 mRNA:sgRNA ratios (0.11 to 39 by weight) at a total RNA dose of 10 ng per well. Gene editing efficiencies exceeded 69% at all tested ratios, with an optimal range between 0.5 and 2, achieving over 85% efficiency. A clear dose-dependent effect was observed, as increasing the total RNA dose to 20 ng per well further improved gene editing performance. Specifically, efficiencies in the 0.5 to 2 ratio range exceeded 92%.

**Figure 7.**
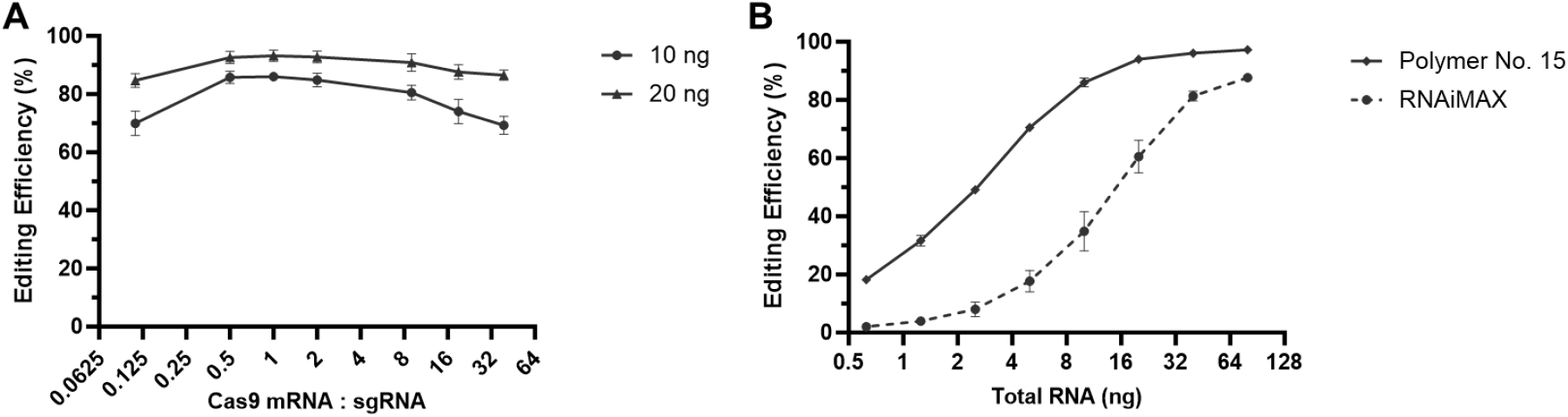
**Polymer No. 15** enables gene-editing techniques in Huh7 cells. (A) Systematic titration of the spCas9 mRNA:sgRNA ratio at a constant total RNA amount of 10 ng or 20 ng. (B) RNA dose titration for spCas9 mRNA and sgRNA at a fixed mRNA:sgRNA ratio of 2:1.

Based on these findings, a spCas9 mRNA:sgRNA ratio of 2 was chosen for the subsequent dose-response study, with the commercially available RNAiMAX serving as the positive control. The total RNA dose varied from 0.625 to 80 ng per well. As shown in **Figure 7B**, gene editing efficiency increased with the total RNA dose, reaching a maximum of 97% at 80 ng. Notably, **Polymer No. 15** outperformed RNAiMAX across all tested doses.

### 6. PBAA random copolymer encapsulates mRNA to form stable nanoparticles

Polyethylene glycol (PEG) is commonly incorporated during formulation to improve the stability of nanoparticles.^[36,37]^ PEGylation of the nanoparticle surface creates a steric barrier that prevents aggregation and increases resistance to ionic strength, thereby enhancing nanoparticle stability.^[38]^ Furthermore, the PEG layer shields the nanoparticle core from interactions with biomolecules, thereby extending circulation time *in vivo*.^[39]^ However, excessive PEG on the nanoparticle surface can hinder nanoparticle-cell interactions and reduce cellular uptake.^[40]^ To investigate the stability of nanoparticles formed with PBAA random copolymers and to optimize formulation conditions that balance nanoparticle stability with RNA delivery efficacy, we prepared nanoparticles using **Polymer No. 15** and a mixture of spCas9 mRNA and sgRNA at a 2:1 weight ratio. DMG-PEG2000 (25 mg/mL in ethanol) was added at volume ratios of 0%, 0.1%, and 1% (% v/v) relative to the total formulation volume, which corresponds to weight ratios of 0%, 5.1%, and 50.9% relative to **Polymer No. 15**. The samples were prepared at an N/P ratio of 10, resulting in a total RNA concentration of 40 ng/µL.

The nanoparticle stabilities over time were first investigated. Formulated solutions were stored at 4 °C for two weeks, with size, PDI, and zeta potential measurements performed at 0 h, 5 h, 1 d, 2 d, 3 d, 7 d, and 14 d using DLS. Prior to measurement, 75 µL of the sample was diluted with water to a total volume of 1000 µL for size and PDI analysis. For zeta potential measurements, 0.05X PBS was used as the diluent instead of water.

As shown in **Figure 8A**, nanoparticles without PEG maintained stable hydrodynamic diameters for one week, while increasing from 200 nm to approximately 280 nm after two weeks of storage. The addition of 0.1% or 1% PEG did not significantly improve size stability, as they followed a similar trend. However, 1% PEG reduced the initial particle size from 200 nm to 160 nm in freshly prepared samples. PEG addition had no impact on sample homogeneity, as the PDI remained consistent across all conditions throughout the two-week storage period (**Figure 8B**). In contrast to size and PDI, the addition of 0.1% PEG significantly reduced the zeta potential from 40 mV to 19 mV, and increasing the PEG concentration to 1% further decreased it to 15 mV (**Figure 8C**). This demonstrates the charge-neutralizing and surface-shielding properties provided by the PEG layer.

**Figure 8.**
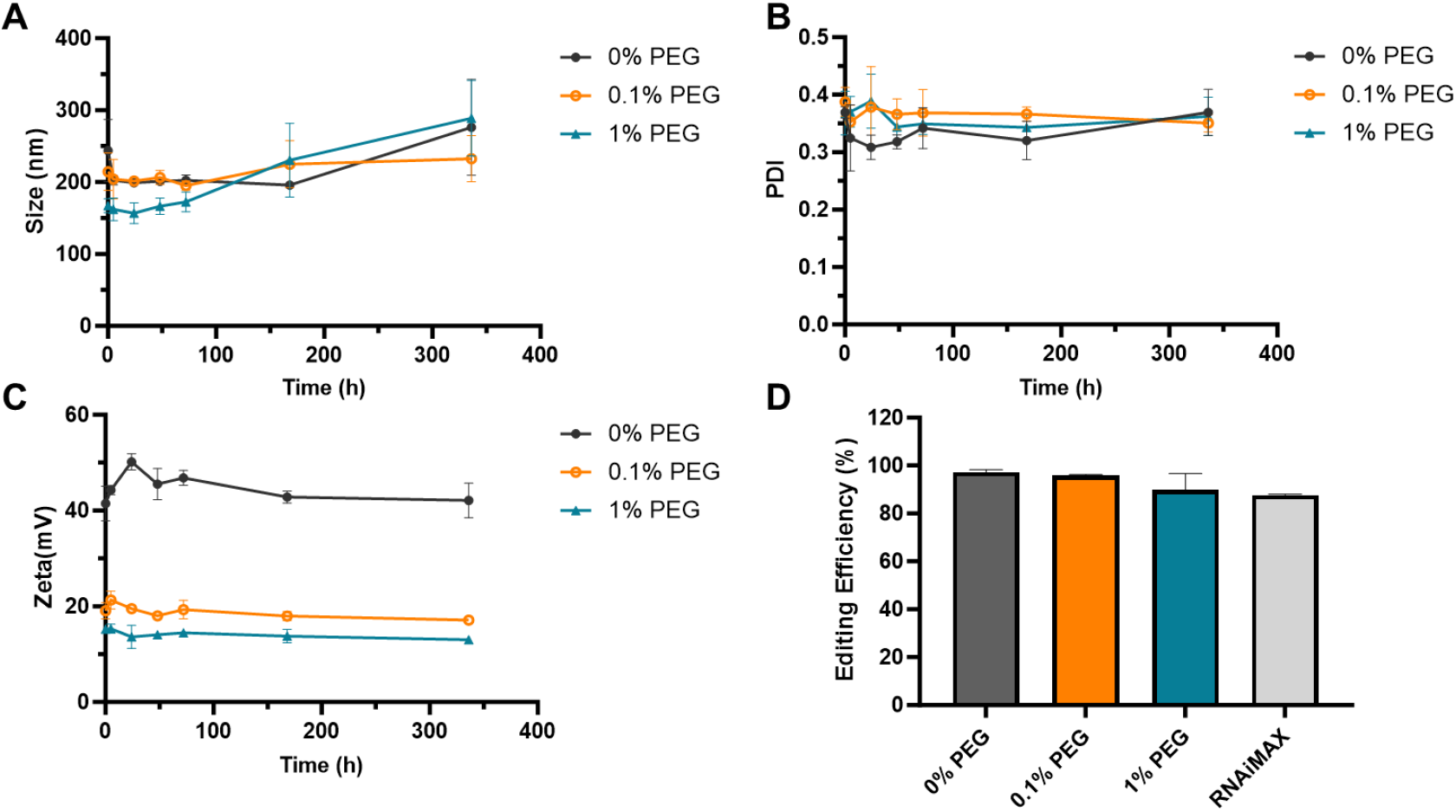
Physicochemical characterization of **Polymer No. 15**/RNA nanoparticles prepared with 0%, 0.1%, and 1% DMG-PEG2000 (expressed as % v/v of a 25 mg/mL DMG-PEG2000 stock solution relative to the total formulation volume). Particle size (A), PDI (B), and Zeta potential (C) were monitored over time at 4 °C. (D) The impact of PEG additives on gene-editing efficiency was assessed in Huh7 cells.

The impact of PEG additives on gene-editing performance was assessed in Huh7 cells. Samples were prepared following the same protocol as the stability test, and cells were treated with a total RNA dose of 80 ng/well. As shown in **Figure 8D**, the formulation containing 0.1% PEG achieved a comparable performance (96%) to the formulation without PEG (97%), whereas 1% PEG led to a more pronounced reduction in performance to 90%. Notably, all tested conditions outperformed the positive control, RNAiMAX, at the same RNA dose.

In summary, **Polymer No. 15** produced stable nanoparticles. Although PEG addition significantly reduced surface charge, its effect on nanoparticle size and PDI was minimal. Among the tested conditions, 0.1% PEG emerged as the optimal concentration, maintaining nanoparticle size and PDI while preserving high gene-editing efficiency. Furthermore, it offers potential benefits for applications that require a reduced zeta potential.

## Conclusion

Functional PBAA polymers have been developed and optimized as a versatile platform for RNA delivery. Through systematic *in vitro* and *in vivo* screening and investigation of SARs, we identified several key findings: (1) incorporating hydrophobic and linear alkyl or alkenyl groups in the side chains and/or chain-end groups significantly outperformed hydrophilic and branched counterparts; (2) increasing the length of alkyl groups enhanced transfection efficacy, with an optimal range between C_12_ and C_18_; and (3) random copolymers with higher lipid content outperformed homopolymers with chain-end modifications, with a lipid content of approximately 0.8 proving optimal for most random copolymers bearing various hydrophobic side chains. The top-performing random copolymers with dodecyl side chains emerged as leading candidates, demonstrating exceptional RNA transfection efficiency, over 97% spleen-targeting selectivity, and more than 97% gene-editing efficacy *in vitro*. These polymers formed stable nanoparticles that maintain their physicochemical properties at 4 °C for up to two weeks without the need for adding PEG. Further formulation optimization, including N/P ratio screening and PEGylation, was conducted to achieve an optimal balance between stability and efficacy. Overall, PBAA polymers, with their capacity for efficient RNA delivery, spleen targeting, and gene editing, offer an adaptable platform for future gene therapy applications.

## Patents

X.Y., J.X., and X.Z. are co-inventors on patents filed by N1 Life, Inc. that cover technologies discussed in the manuscript.

## Data Availability Statement

The data presented in this study are available on request from the corresponding author.

## Acknowledgments

The authors thank Dr. Jiuzhi Sun for performing flow cytometry, and Glow Biosciences LLC for performing *in vivo* studies. Special thanks to Dr. Jiuzhi Sun for the discussion.

