## Supplementary Material for "Biodegradable Poly(*β*-amino amide)s Enable Efficient RNA Delivery and Spleen Targeting"

Table of Contents

1. Chemicals, Reagents, and Instruments .......... S2

2. Polymer Synthesis .......... S3

2.1 General Synthetic Procedure for Homopolymer .......... S3

2.2 General Synthetic Procedure of Polymers with Chain End Modification .......... S3

2.3 General Synthetic Procedure for Random Copolymer .......... S4

2.4 General Synthetic Procedure for Polymer Deprotection .......... S5

3. Polymer/RNA complex preparation .......... S5

3.1 General Formulation Method 1- Pipetting Method .......... S5

3.2 General Formulation Method 2- Shaker Method .......... S5

4. *In Vitro* mRNA Transfection .......... S6

4.1 Cell Culture .......... S6

4.2 In Vitro eGFP mRNA Transfection .......... S6

4.3 In Vitro Fluc mRNA Transfection .......... S7

5. *In Vitro* Gene Editing .......... S7

5.1 Cell Culture .......... S7

5.2 In Vitro Gene Editing Formulation .......... S7

5.3 In Vitro Gene Editing Efficiency Determination .......... S8

6. *In Vivo* Fluc mRNA Transfection .......... S8

6.1 Animal .......... S8

6.2 In Vivo Polymer/Fluc mRNA Formulation .......... S8

6.3 BLI Imaging .......... S9

6.4 Analytic Method .......... S9

7. Nanoparticle Stability Study .......... S10

Figure S1.. .......... S10

Table S1.. .......... S11

### 1. Chemicals, Reagents, and Instruments

All chemicals and solvents were acquired from commercial sources (Sigma-Aldrich, VWR, Fisher, TCI, Oakwood Chemicals, ChemScene) and used without purification unless otherwise noted. All consumables such as syringes, needles, syringe filters, glass vials, etc. were acquired from commercial sources (VWR, Fisher, Zoro, Chemglass).

All reagents used for the in vitro and in vivo assays were acquired from commercial sources as listed below: Dulbecco's Modified Eagle Medium (DMEM) (10599010, Gibco), Opti-MEM (31985062, Gibco), 0.05% Trypsin-EDTA (25300054, Gibco), Certified Fetal Bovine Serum (C04001-500, VivaCell), PBS pH 7.4 (1×) (10010-049, Gibco), penicillin-streptomycin solution (C0222, Beyotime), Lipofectamine 3000 (L3000-001, Invitrogen), Lipo Messenger Max (LMRNA015, Thermo), UltraPure™ DNase/RNase-Free Distilled Water(10977015, Thermo), 0.4% Trypan blue (15250061, Thermo), human fibronectin (FC010, Sigma-Aldrich), IVISbrite D-Luciferin potassium salt (122799, PerkinElmer), DMG-PEG2000 (C08501-N230201, Jenkem), eGFP mRNA (L-7601, Trilink), Fluc mRNA (L-7602, Trilink), spCas9 mRNA and sgRNA-2401 (were synthesized by AccurEdit-Therapeutics)

$^{1}$H NMR was recorded on Varian 400, 500, or 600 MHz spectrometers. Chemical shifts are reported in delta ($\delta$) units, expressed in parts per million (ppm) downfield from tetramethylsilane using protio-solvent (residual) as internal standard ($CD_3OD$, $\delta$H = 3.31 ppm for $^{1}$H NMR, $D_2O$, $\delta$H = 4.79 ppm for $^{1}$H NMR). Data are reported as chemical shift ($\delta$), multiplicity (s = singlet, d = doublet, t = triplet, m = multiplet, br = broad), and integration.

Instruments used in this study included Varioskan® Flash (3001-1375, Thermo Scientific), EVOS Imaging System (M5000, ThermoFisher Scientific), Microplate Reader (Infinite M200 PRO, TECAN), Inverted Routine Microscope (TS100, Nikon), IVIS Lumina X5 imager (Perkin Elmer), and Living Image Software (PerkinElmer), Panalytical Zetasizer Ultra SystemZetasizer Ultra (ZSU3305, Malvern Panalytical), Intelligent Cell Analyzer (Rigel S5, Count Star).

### 2. Polymer Synthesis

*2.1 General Synthetic Procedure for Homopolymer*

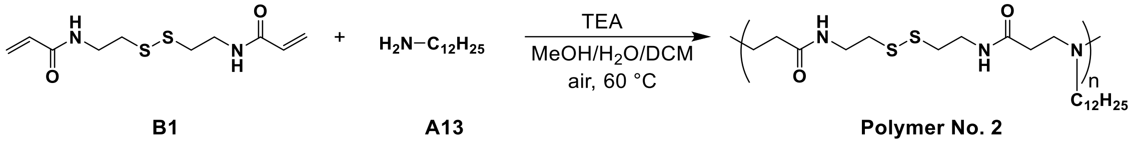


The homopolymers were synthesized following a two-step procedure. First, the polymer backbone was synthesized by Michael addition of a primary amine with a diacrylate at a 1:1 mole ratio (amine:diacrylate). Subsequently, additional amine used in the first step was added to cap any free acrylate chain ends. The general procedure for synthesizing homopolymers is described here, using **Polymer No. 2** as an example: N,N′-Bis(acryloyl)cystamine (**B1**, 1 eq.), dodecylamine (**A13**, 1 eq.), and triethylamine (0.1 eq.) were added into a vial charged with a stir bar. The above reagents were suspended in a MeOH/water/DCM mixture (4/1/2.1, v/v/v) ($[\mathbf{B1}]_0$ = 0.51 M) and the vial was sealed with a PTFE-lined screw cap under air. The reaction was stirred at 60 °C for 48 h. Upon cooling down, extra dodecylamine (**A13**, 0.2 eq.) in a DCM solution (0.48 M) was added to the above mixture. The vial was sealed with a PTFE-lined screw cap under air. The reaction was continued at 60 °C for 24 h. Upon completion, the reaction was cooled down to room temperature. The crude polymer sample was diluted with an equal volume of DCM/MeOH (1:1, v/v) and subsequently precipitated into cold hexane at 30 times its volume. The polymer was collected by centrifugation at 3000 rpm under -10 °C for 5 min and washed with cold hexane three times. After repeated the precipitation process, the polymer was collected and dried under vacuum, giving **Polymer No. 2** as a white gooey solid.

*2.2 General Synthetic Procedure of Polymers with Chain End Modification*

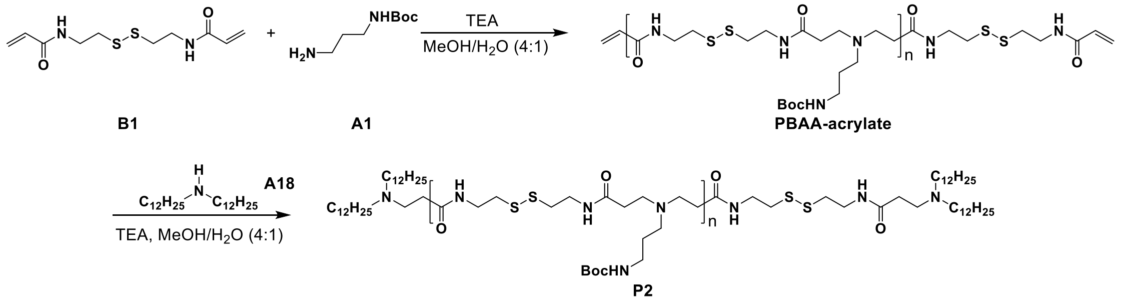


The chain-end-modified polymers were synthesized following a two-step procedure. First, an acrylate-terminated polymer was synthesized by Michael addition of primary amines with diacrylates at a 1:1.2 mole ratio (amine:diacrylate). Next, the acrylate chain ends were functionalized with various primary or secondary amines. The general procedure for synthesizing chain end modified homopolymers is described here, using polymer **P2** (the Boc-protected precursor of **Polymer No. 8**) as an example: N, N′-

Bis(acryloyl)cystamine (**B1**, 1.2 eq.), N-Boc-1,3-propanediamine (**A1**, 1 eq.), and triethylamine (0.1 eq.) were added into a vial charged with a stir bar. The above reagents were suspended in a MeOH/water mixture (4/1, v/v) ([**B1**]$_0$ = 0.60 M) and the vial was sealed with a PTFE-lined screw cap under air. The reaction was stirred at 60 °C for 48 h. Upon completion, the reaction was cooled down to room temperature. The crude polymer sample was diluted with an equal volume of DCM/MeOH (1:1, v/v) and subsequently precipitated into cold $Et_2O$ at 30 times its volume. The polymer was collected by centrifugation at 3000 rpm under -10 °C for 5 min, washed with cold $Et_2O$ three times, and dried under vacuum, giving **PBAA-acrylate** as a white solid.

In the second step of chain-end modification, suspended the **PBAA-acrylate** polymer in a MeOH/water solvent mixture (5/1, v/v) (250 mg/mL). Then didodecylamine solution (**A18**, 0.4 eq.) in DCM (0.43 M) and triethylamine (0.1 eq.) were added consecutively. The vial was sealed with a PTFE-lined screw cap under air. The reaction was stirred at 60 °C for 24 h. Upon completion, the reaction was cooled down to room temperature. The crude polymer sample was diluted with an equal volume of DCM/MeOH (3/1, v/v) and subsequently precipitated into cold hexane at 30 times its volume. The polymer was collected by centrifugation at 3000 rpm under -10 °C for 5 min, washed with cold hexane three times, and dried under vacuum, giving **Polymer No.8** as a white solid.

*2.3 General Synthetic Procedure for Random Copolymer*

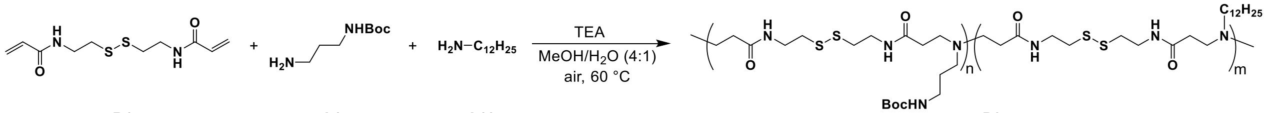


B1 A1 A13 P3

The random copolymers were synthesized as described below, using polymer **P3** (the Boc-protected precursor precursor of **Polymer No. 15**) as an example: N,N′-Bis(acryloyl)cystamine (**B1**, 1 eq.), N-Boc-1,3-propanediamine (**A1**), dodecylamine (**A13**) (**A1** + **A13** = 1 eq.) and triethylamine (0.1 eq.) were added into a vial charged with a stir bar. The above reagents were suspended in a MeOH/water/DCM mixture (4/1/2.1, v/v/v) ([**B1**]$_0$ = 0.60 M) and the vial was sealed with a PTFE-lined screw cap under air. The reaction was stirred at 60 °C for 48 h. Upon cooling down, extra dodecylamine (**A13**, 0.10 eq.) in a DCM solution (0.35 M) was added into the above mixture. The vial was sealed with a PTFE-lined screw cap under air. The reaction continued at 60 °C for 24 h. Upon completion, the reaction was cooled down to room temperature. The crude polymer sample was diluted with an equal volume of DCM/MeOH (2/1, v/v) and subsequently precipitated into cold hexane at 30 times its volume. The polymer was collected by centrifugation at 3000 rpm under -10 °C for 5 min and washed with cold hexane three times. After repeated the precipitation process, the polymer was collected and dried under vacuum, giving **P3** as a white solid.

*2.4 General Synthetic Procedure for Polymer Deprotection*

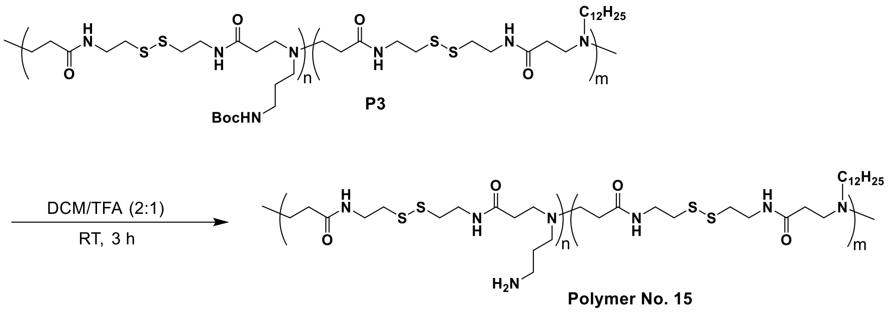


The general procedure for the deprotection of -NHBoc is detailed below, with the synthesis of **Polymer No. 15** as an example: To a vial containing polymer P3 and a stir bar was added a mixture of degassed DCM and trifluoroacetic acid (2/1, v/v), achieving a concentration of 20 mg/mL. The reaction was sealed under an inert atmosphere and stirred at room temperature for 3 h. After completion, the reaction mixture was concentrated and dried under vacuum, yielding the deprotected polymer (**Polymer No. 15**).

### 3. Polymer/RNA complex preparation

*3.1 General Formulation Method 1- Pipetting Method*

Polymer/RNA complex solutions were prepared with polymer-to-RNA molar ratios (N/P ratios) ranging from 5:1 to 25:1. In a 1.5 mL microcentrifuge tube, the specified amount of RNase-free water was added, followed by the RNA stock solution (0.1 or 1 mg/mL in RNase-free water or 1X PBS buffer) at room temperature. The solution was briefly mixed by pipetting. The RNA used was either a single component or a premixed blend. Next, varying amounts of polymer stock solution (2.5 or 5 mg/mL in RNase-free water) were added. The solution was thoroughly mixed by pipetting 20 times (about 1 time/s). For groups with PEG additives, a solution of DMG-PEG2000 (25 mg/mL in EtOH) was added after the polymer stock and mixed by pipetting 10 times. All samples were incubated at room temperature for 5 min.

*3.2 General Formulation Method 2- Shaker Method*

In certain studies, polymer/RNA complexes were prepared using a shaker. The polymer stock solution (2.5 or 5 mg/mL in RNase-free water) was first diluted in water, followed by the addition of the RNA stock solution (0.1 or 1 mg/mL in RNase-free water or 1X PBS buffer). The RNA consisted of either a single component or a premixed blend. The solution was briefly mixed by pipetting 3 times (about 1 time/s) at each step, then further mixed on a shaker at 900 rpm for 30 s at 25 °C. The sample was subsequently incubated at room temperature for 5 min. For groups with PEG additives, a solution of DMG-PEG2000 (25 mg/mL in EtOH) was added after the polymer stock and mixed by pipetting 3 times (about 1 time/s) before the RNA stock solution was introduced.

### 4. *In Vitro* mRNA Transfection

#### *4.1 Cell Culture*

HEK 293T cells were cultured in Dulbecco's Modified Eagle Medium (DMEM) supplemented with 10% (v/v) fetal bovine serum (FBS) and 1% (v/v) penicillin-streptomycin. The cells were maintained at 37°C in a humidified atmosphere with 5% $CO_2$. For cell passaging, 0.05% trypsin-EDTA was used to dissociate the adherent cells.

#### *4.2 In Vitro eGFP mRNA Transfection*

HEK 293T cells were seeded at 10,000 cells/well in 100 µL DMEM supplemented with 10% (v/v) FBS and 1% (v/v) penicillin-streptomycin in black-walled 96-well plates pre-coated with human fibronectin. Cells were incubated 18–24 h at 37 ˚C (5% $CO_2$) prior to transfection ensuring that confluency reaches 60%–80%. Right before transfection, the serum-containing media in each well were removed, and the cells were washed once with 100 µL of serum-free DMEM media per well. Then 50 µL of Opti-MEM was added to each well.

Polymer/eGFP mRNA polyplexes were prepared at an N/P ratio of 25 with an mRNA dose of 40 ng per well following the general formulation method 1. For example, to prepare **Polymer No. 15**/mRNA polyplexes, 24.9 µL of RNase-free water was added to a 1.5 mL microcentrifuge tube, followed by 3.20 µL of mRNA stock solution (0.1 mg/mL in 1X PBS buffer). The mixture was pipetted five times to achieve a final mRNA concentration of 0.01 mg/mL. Subsequently, 3.93 µL of **Polymer No. 15** stock solution (2.5 mg/mL in RNase-free water) was added, and the solution was thoroughly mixed by pipetting 20 times (approximately once per second). After incubation at room temperature for 5 min, the sample was diluted with 368 µL of Opti-MEM. A volume of 50 µL of the diluted sample was dispensed into each well, achieving a total volume of 100 µL per well for treatment. The commercially available transfection reagents, such as Lipofectamine™ 2000 and jetMESSENGER, were used as positive controls and prepared according to the manufacturer's instructions. Six replicates were run for each condition. After 4 h of incubation at 37 °C, 100 µL of complete media was added to each well. The transfection efficacy was assessed 24 h post-transfection using the EVOS M500 Imaging System.

For the lipid content screening *in vitro*, flow cytometry was performed 4 h and 28 h post-transfection to evaluate transfection efficiency and eGFP expression levels. The data are presented as the average percentage of GFP-positive cells and mean fluorescence intensities (MFI), with error expressed as ± SD.

*4.3 In Vitro Fluc mRNA Transfection*

The studies were conducted using HEK 293T cells. Polymer/Fluc mRNA polyplexes preparation follows the same procedure as described in Section 4.2, with a mRNA dose of 40 ng/well and a N/P ratio of 25. Lipofectamine™ 2000 was used as a positive control and prepared according to the manufacturer's instructions. Six replicates were run for each condition. After 4 h of incubation at 37 °C with 5% $CO_2$, 100 µL of complete media was added to each well. At 24 h post-transfection, 10 µL of D-luciferin solution (1.5 mg/mL in Opti-MEM) was added to each well. The plate was shaken for 5 s using a plate reader and then incubated at 37 °C with 5% $CO_2$ for 5 minutes. Bioluminescent signals were recorded using the TECAN Infinite M200 PRO plate reader. Data are presented as mean bioluminescence intensities, with errors expressed as ± SD.

### 5. *In Vitro* Gene Editing

*5.1 Cell Culture*

Huh7 cells were cultured in Dulbecco's Modified Eagle Medium (DMEM) supplemented with 10% (v/v) fetal bovine serum (FBS). The cells were maintained at 37°C in a humidified atmosphere with 5% $CO_2$. For cell passaging, 0.05% trypsin-EDTA was used to dissociate the adherent cells. Cells were seeded at a density of 8,000 cells per well in 100 µL of DMEM in a 96-well plate and incubated for 24 hours prior to transfection.

*5.2 In Vitro Gene Editing Formulation*

For the spCas9 mRNA:sgRNA ratio titration experiments, spCas9 mRNA and sgRNA (targeting the genomic sequence of AAAGGCTGCTGAUGACACCT) were prepared as 0.1 mg/mL stock solutions. Then, spCas9 mRNA and sgRNA were premixed at various weight ratios ranging from 0.11 to 39. **Polymer No. 15**/RNA polyplexes were prepared following the general formulation method 2. For example, 87.5 µL of RNase-free water was added to a 1.5 mL microcentrifuge tube, followed by 2.5 µL of **Polymer No. 15** stock solution (5 mg/mL in RNase-free water). Then 10 µL of the RNA mixture was added. The solution was briefly mixed by pipetting 3 times (about 1 time/s) at each step, then further mixed on a shaker at 900 rpm for 30 s at 25 °C, achieving a final RNA concentration of 10 ng/µL. After incubation at room temperature for 5 min, the sample was added to each well to achieve a total RNA dose of 10 ng or 20 ng and a final volume of 100 µL per well.

For the RNA dose titration experiment, spCas9 mRNA and sgRNA (targeting the genomic sequence of AAAGGCTGCTGAUGACACCT) were prepared as 1 mg/mL stock solutions. Then, the spCas9 mRNA and sgRNA were premixed at a fixed ratio of 2:1. **Polymer No. 15**/RNA polyplexes were prepared using the same method described above (general formulation method 2). The total RNA dose varied from 0.625 to 80 ng per well.

#### *5.3 In Vitro Gene Editing Efficiency Determination*

72 h after transfection, genomic DNA was extracted using QuickExtract DNA Extraction Solution (Lucigen #QE09050) per manufacturer's protocol. Amplicon libraries were generated by PCR amplification of target regions using locus-specific primers (F: CGCTCCAGATTTCTAATACCACA, R: GTCCTGTGGGAGGGTTCTTT). PCR reactions were performed using Q5 Hot Start HiFi PCR Master Mix (NEB #M0543L). Thermocycling conditions were as follows: initial denaturation at 98℃ for 5 min, followed by 35 cycles of denaturation at 98 ℃ for 30 s, annealing at 60 ℃ for 90 s, and extension at 72 ℃ for 30 s, with a final extension at 72 ℃ for 10 min. Then second round PCR amplification was performed to add illumina adapter sequences. PCR products were purified using VAHTS DNA Clean Beads (Vazyme # N411-02) and sequenced on an Illumina NovaSeq 6000 platform with paired-end 150 bp according to the manufacturer’s instructions.

The sequencing data was subjected to quality control and paired-end merging using fastp (doi.org/10.1093/bioinformatics/bty560). Then the quality-controlled data was analyzed using an enhanced version of CRISPResso2(doi.org/10.1038/s41587-019-0032-3) (quantification_window_size 3) to obtain the editing efficiency data of sgRNA.

### 6. *In Vivo* Fluc mRNA Transfection

#### *6.1 Animal*

All animal experiments were consistent with local, state, and federal regulations as applicable. Food and water were supplied *ad libitum*. Female BALB/c mice (Charles River, Cambridge, UK), and female ICR mice, aged 6−8 weeks, were randomly assigned to groups and housed in a fully climate-controlled room.

#### *6.2 In Vivo Polymer/Fluc mRNA Formulation*

Polymer/Fluc mRNA (Trilink L-7602) polyplexes were formulated using general formulation method 1. For example, to prepare **Polymer No. 15**/mRNA polyplexes for the first and second round of study, 579 µL RNase-free water was added to a 1.5 mL microcentrifuge tube, followed by 20 µL Fluc mRNA (1 mg/mL solution in 1X PBS). The solution was mixed by pipetting 10 times, followed by the addition of 121 µL **Polymer No.15** (5 mg/mL stock solution in RNase-free water). The sample was further mixed by pipetting 30 times (about 1 time/s). Then, 80 µL 10X PBS was added, giving a total volume of 800 µL with a 1X PBS concentration. The solution was thoroughly mixed, and 200 µL of the mixture, containing 5 µg of Fluc mRNA per mouse, was injected intravenously into the tail vein of female BALB/c mice. The negative control group, containing only 5 µg of Fluc mRNA per mouse, was prepared following the same method.

*6.3 BLI Imaging*

The biodistribution of polymer/Fluc mRNA polyplexes in mice was assessed using the IVIS Lumina X5 imaging system (PerkinElmer). A fresh luciferin stock solution (30 mg/mL in 1X PBS) was prepared, and each mouse received an intraperitoneal injection of 100 µL of the solution 10 minutes prior to imaging. Mice were anesthetized with isoflurane and imaged using Living Image software (PerkinElmer) with automatic imaging settings (exposure time ranging from 20 s to 4 min, depending on signal strength). Whole body images were captured in both ventral and lateral positions at each time point.

In the first-round *in vivo* study, female BALB/c mice were imaged at 6 and 16 h post-administration. After the 16 h imaging, mice were immediately sacrificed. Major organs, including the brain, heart, lungs, liver, spleen, stomach, intestines, kidneys, and skeletal muscle, were excised and imaged *ex vivo* from two mice per group.

In the second-round *in vivo* study, female BALB/c mice were imaged at 3, 6, and 12 h post-administration. Two mice were sacrificed at the 6 h time point, and one at the 12 h time point. Major organs, including the brain, heart, lungs, liver, spleen, stomach, intestines, kidneys, and skeletal muscle, were promptly harvested and imaged *ex vivo*.

In the third-round *in vivo* study, female ICR mice were imaged at 6 h post-administration, followed by immediate sacrifice. Major organs, including the spleen, liver, and lungs, were promptly harvested and imaged *ex vivo*.

*6.4 Analytic Method*

Living Image software was utilized to analyze the numerical pixel values and image data from the acquisition files for all experimental groups. The saved Living Image files were loaded for image display and analysis. Circular Regions of Interest (ROIs) were placed around each luminescent source, and the system automatically displayed data for all the ROIs created within the images.

### 7. Nanoparticle Stability Study

The samples used for stability test were prepared using **Polymer No. 15** and a mixture of spCas9 mRNA and sgRNA (1 mg/mL in RNase-free water) at a weight ratio of 2:1. DMG-PEG2000 (25 mg/mL in EtOH) was added at volume ratios of 0%, 0.1%, and 1% (% v/v) relative to the total formulation volume. Polymer/RNA polyplexes were prepared following the general formulation method 2, with an N/P ratio of 10. For example, to prepare **Polymer No. 15**/RNA polyplexes with 0.1% PEG, 1291.2 µL of RNase-free water was added to a 2 mL microcentrifuge tube, followed by 147.3 µL of **Polymer No. 15** stock solution (5 mg/mL in RNase-free water) and 1.5 µL of DMG-PEG2000 (25 mg/mL in EtOH). Next, 60 µL of the RNA mixture was added. The solution was briefly mixed by pipetting 3 times (about 1 time/s) at each step, then further mixed on a shaker at 900 rpm for 30 s at 25 °C, achieving a final RNA concentration of 40 ng/µL. After incubation at room temperature for 5 min, the samples were stored at 4 °C for two weeks. Size, PDI, and zeta potential measurements were performed at 0 h, 5 h, 1 d, 2 d, 3 d, 7 d, and 14 d using Malvern Panalytical Zetasizer Ultra System. Prior to measurement, 75 µL of the sample was diluted with water to a total volume of 1000 µL for size and PDI analysis. For zeta potential measurements, 0.05X PBS was used as the diluent instead of water.

| Polymer No. | $NH_2/C_{12}$ |
|---|---|
| 17 | 4.68 |
| 18 | 3.02 |
| 19 | 1.82 |
| 20 | 0.78 |
| 21 | 0.31 |
| 22 | 0.14 |
| 23 | 0.04 |

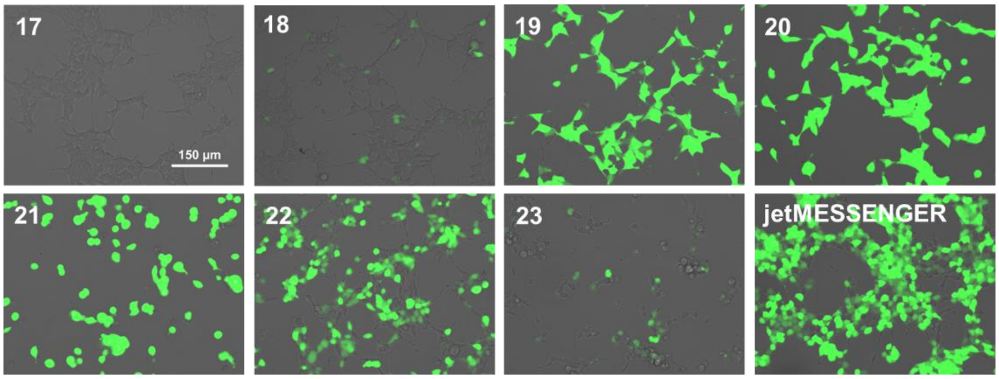


**Figure S1.** Fluorescence microscopy images of HEK 293T cells after 24 hours of treatment with PBAA/eGFP mRNA nanocomplex were captured to assess the performance of dodecyl random copolymers **No. 17** to **23** (Scale bar, 150 µm). The lipid contents of these copolymers are shown in the table on the left.

**Table S1.** Experimental details for each group in the third-round *in vivo* study on Fluc mRNA delivery.

| Group | Test article | N/P | mRNA dose (µg per mouse) | Route | Animal number |
|---|---|---|---|---|---|
| 1 | Vehicle | / | / | i.v. | 1♀ |
| 2 | Polymer No. 15 | 5 | 50 | i.v. | 2♀ |
| 3 | Polymer No. 15 | 10 | 25 | i.v. | 2♀ |
| 4 | Polymer No. 15 | 25 | 10 | i.v. | 2♀ |
| 5 | Polymer No. 16 | 5 | 50 | i.v. | 2♀ |
| 6 | Polymer No. 16 | 10 | 25 | i.v. | 2♀ |
| 7 | Polymer No. 16 | 25 | 10 | i.v. | 2♀ |
